# Altered Striatal Acetylcholine Dynamics in a Model of Parkinson’s Disease and L-DOPA-induced Dyskinesia

**DOI:** 10.64898/2026.06.11.731672

**Authors:** Sophie C Gregoretti, Karen L Eskow Jaunarajs, Mariangela Scarduzio

**Affiliations:** Killion Center for Neurodegeneration and Experimental Therapeutics; Department of Neurology, UAB, Birmingham, AL, USA

**Author notes:** Corresponding Author: Mariangela Scarduzio, PhD, Center for Neurodegeneration and Experimental Therapeutics, Department of Neurology, UAB| The University of Alabama at Birmingham CIRC 345B| 1719 6th Ave S| Birmingham, AL 35294.

**Keywords:** striatum, movement disorders, acetylcholine, PD, L-DOPA-induced dyskinesia, fiber photometry, cholinergic interneurons

## Abstract

L-DOPA remains the most effective therapy for Parkinson’s disease (PD), yet its chronic use often induces involuntary movements known as L-DOPA–induced dyskinesia (LID). While abnormal cholinergic interneuron (ChI) activity is a hallmark of both PD and LID, emerging evidence suggests that the temporal organization of acetylcholine (ACh) signaling, rather than its overall magnitude, may determine its functional impact. Under physiological conditions, ChIs exhibit intrinsic delta-frequency activity reflected in coordinated, slow oscillations of extracellular ACh, which are thought to organize striatal network function and movement pattering.

To determine how dopamine (DA) depletion and L-DOPA treatment reshape these ACh dynamics, we used *in vivo* GRAB-ACh fiber photometry in the unilateral 6-OHDA mouse model of parkinsonism. DA depletion disrupted slow ACh rhythmicity, reducing delta-band regularity while increasing higher-frequency phasic activity. Acute L-DOPA broadly suppressed ACh activity across frequencies, partially normalizing this imbalance, but without restoring slow temporal structure. In addition, chronic L-DOPA treatment, associated with established dyskinesia, further impaired delta-band coordination in the DA-depleted striatum during the ON state, while OFF-state activity retained lesion-associated features. The anti-dyskinetic agent amantadine restored low-frequency temporal structure both before and after L-DOPA exposure.

Together, these findings reveal a state-dependent reorganization of striatal ACh dynamics, characterized by a shift from coordinated slow oscillations to irregular phasic activity following DA loss, and a further breakdown of slow temporal organization during dyskinetic states. These results highlight the temporal structure of cholinergic signaling as a critical and underappreciated dimension of striatal function in PD and LID.

## INTRODUCTION

Dopamine (DA) and acetylcholine (ACh) interact closely within the dorsal striatum to regulate motor control and action selection. Classical models of basal ganglia function emphasize a reciprocal interaction between DA and ACh, in which the relative balance of these neuromodulators shapes striatal output and motor function ^[1, 2]^. In Parkinson’s disease (PD), loss of nigrostriatal DA and its pharmacological replacement with L-DOPA are both associated with profound alterations in motor behavior and prevailing frameworks have largely interpreted these effects through changes in DA and ACh levels or their receptor activation ^[3–6]^. Within this view, motor abnormalities and L-DOPA–induced dyskinesia (LID) are commonly linked to disrupted DA–ACh balance ^[7]^. Recent methodological advances, especially genetically encoded fluorescent sensors allowing direct monitoring of neuromodulator dynamics in vivo ^[8]^, have begun to challenge this static perspective by revealing that, in the striatum, both DA and ACh signals exhibit spontaneous fluctuations rather than a static tonic activity ^[9]^.

Under physiological conditions, cholinergic interneurons (ChIs) display intrinsic membrane oscillations at delta frequencies (1–4 Hz) ^[10]^ and synchronized firing patterns that give rise to rhythmic fluctuations in extracellular ACh ^[9, 11]^. ChI occupy a unique position within striatal circuits. Despite their sparse numbers, they exert widespread influence over corticostriatal transmission, DA release, local interneuron activity, and the excitability of both direct- and indirect-pathway spiny projection neurons. Through these actions, ChIs are well positioned to coordinate activity across large neuronal populations. Indeed, ACh rhythmicity has been associated with coordination of striatal network oscillatory activity and movement patterning ^[10]^ and disruptions of the ACh rhythm were found in association with movement abnormalities in a paroxysmal genetic model of dyskinesia/dystonia ^[11]^. These observations suggest that ACh neuromodulatory signaling carries information not only through magnitude, but also through temporal structure, extending traditional balance models toward a dynamic framework.

Both suppression ^[12–14]^ and enhancement of cholinergic signaling ^[15]^ can alleviate dyskinesia, and optogenetic activation of ChIs can either mitigate or exacerbate LID depending on the stimulation patterns used ^[16]^. These observations highlight the temporal structure of cholinergic transmission as a critical, but underexplored, dimension of striatal function, pointing to the pattern rather than the amount of ACh release in rendering striatal circuits susceptible to LID. Nonetheless, precisely how PD and LID-associated DA perturbations reshape the rhythmic structure of striatal ACh transmission in vivo has not been systematically examined. Addressing this gap is critical for understanding how neuromodulatory dynamics are altered across DA-depleted and treated states, and for interpreting the network consequences of pharmacological interventions.

In this study, we tackled this question by investigating how DA loss and L-DOPA treatment affect the oscillatory activity of striatal ACh signaling *in vivo* in the unilateral 6-hydroxydopamine (6-OHDA) mouse model of hemiparkinsonism. We engaged GRAB-ACh fiber photometry to monitoring striatal ACh rhythms across the continuum timeline from the DA lesion, to acute L-DOPA exposure, to the ON and OFF states of chronic L-DOPA treatment. By focusing on oscillatory structure, rather than on single events, we aim to define how distinct dopaminergic states are associated with specific alterations in cholinergic dynamics at the network level. Such knowledge refinement has the potential to improve pharmacological interventions and guide adaptive stimulation protocols to alleviate movement disabilities in PD and LID.

## MATERIALS AND METHODS

All experimental procedures were performed per the National Institutes of Health Guide for the Care and Use of Laboratory Animals with prior review and approval by the University of Alabama Institutional Animal Care and Use Committee (APN: IACUC-22954).

### Animal Model and Experimental Design

The well-established 6-hydroxydopamine (6-OHDA)- induced hemiparkinsonian mouse model was used in this study. Male and female C57Bl/6J mice (20-30 weeks old) underwent unilateral medial forebrain bundle (MFB, from Bregma (mm) AP:-0.7; ML:-1.2; DV:-4.7) 6-OHDA or SHAM (saline) injections (0.2 uL at 3 ug/uL) using stereotaxic methods as previously described ^[17]^. This model induces >90% DA depletion within the ipsilateral striatum and enables the induction of LID upon repeated treatment with L-DOPA. As is traditional with LID animal models, the contralateral hemisphere served as a within-subject not-lesioned control and the SHAM-lesioned animals, which received a dummy injection, as in between-subject controls (**Figure 1A-B**). During the same procedure, bilateral intrastriatal injections (AP:+1; ML:+/- 1.8; DV:-3.2) of fluorescent sensors for ACh were performed (AAV9-hSyn-GRABgACh3.0, 1 uL volume at a titer 1x10e12 vg/mL, Addgene). Fiberoptic probes (200 μm in diameter, 0.37NA, 4mm long, Doric Lenses) were implanted immediately after virus injection at each injection site (**Figure 1C**). Four weeks post-surgery, all mice received one injection of vehicle (0.9% saline, 0.2% ascorbic acid, ip) before starting L-DOPA treatment (1 mg/kg + benserazide, 15 mg/kg, ip) for 10 consecutive days. This dosage was chosen as we have observed a slower induction of LID as compared to higher doses of L-DOPA (2-6 mg/kg), which induce severe LID from the first exposure. Each animal underwent 4 imaging/motor behavior sessions: one on the first day of vehicle injection + on days 1, 5 and 10 of L-DOPA treatment. Each session consisted in 10 min baseline (OFF L-DOPA) followed by 50 min post-injection (ON L-DOPA) for a total of 60 min sessions. Pre-vehicle recordings were used to estimate ACh oscillations between DA-lesioned vs non-lesioned hemispheres and compared to sham-lesioned mice. Within each animal, recording sessions during L-DOPA treatment accounted for both immediate and prolonged effects of L-DOPA exposure and LID development on striatal ACh transmission dynamics. Saline injections were used to control for effects of the injection itself on ACh dynamics. Furthermore, in each imaging session, comparisons between ACh oscillations at baseline versus after L-DOPA injections overtime allowed discerning changes between the OFF and ON states, respectively (**Figure 1D**).

**Figure 1.**
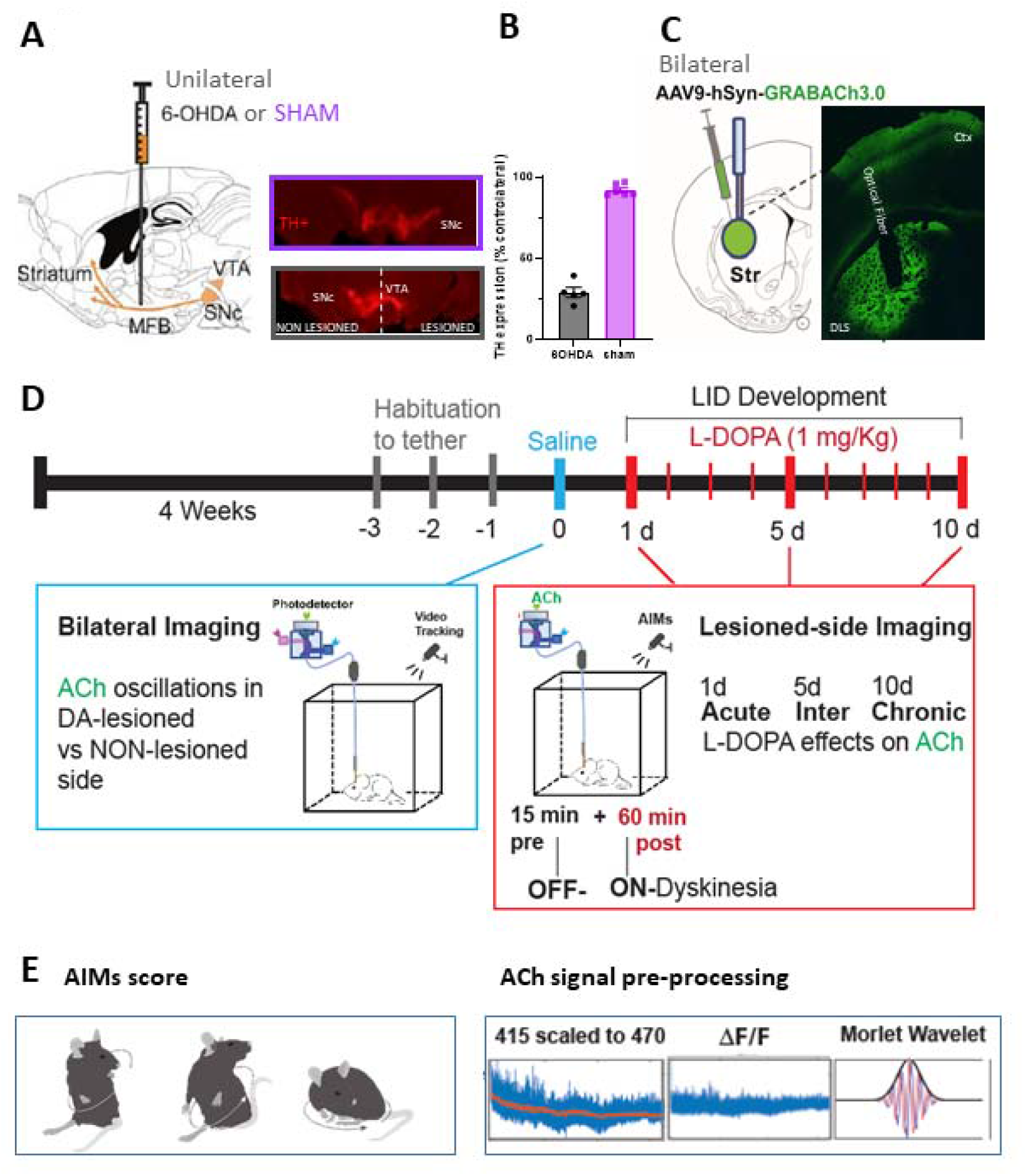
Experimental design. **(A)** Unilateral 6-OHDA injections into the medial forebrain bundle (MFB) produced **(B)** ∼70% depletion of tyrosine hydroxylase immunostained neurons (TH+) in ipsilateral substantia nigra pars compacta (SNc). **(C)** Representative image and schematic depicting expression of the GRAB-ACh sensor and placement of fiber-optic probes in the dorsolateral striatum (DLS) at the site of viral expression. **(D)** Timeline of imaging sessions across the lesion and L-DOPA treatment phases. **(E)** Overview of behavioral assessments and corresponding ACh photometry signal outputs acquired throughout the experimental protocol

### Fiber Photometry

Photometry recordings were performed in awake, freely moving animals in open field via a fiber optic patch cord (200 μm diameter, Doric) coupled to a rotary swivel to permit unrestricted movement. Photometry signals were acquired with the FP3002 system (NPM) controlled via Bonsai software (Bonsai Foundation CIC), which also triggered the video camera. Excitation light was delivered at GRAB-ACh sensor-specific (470nm) and isosbestic (415nm) wavelengths and emitted fluorescence was collected through the same optical fiber. Both LEDs (light power adjusted to 50–100 μW) were alternatively turned on at 100 Hz.

### Photometry Data Analysis

Photometry data were processed using custom scripts in MATLAB (R2023b, MathWorks). Raw fluorescence signals were deinterleaved offline and corrected for motion artifacts, photobleaching drift, and nonspecific fluctuations by fitting the isosbestic reference signal to the ACh-sensitive channel, followed by calculation of ΔF/F.

*Time–frequency representations* of the ΔF/F traces were obtained using continuous Morlet wavelet transforms (MATLAB cwt function; 25 voices per octave) to estimate spectral power between 0.5 and 20⍰Hz (**Figure 1E**). To normalize signal amplitude across animals, reduce 1/f structure and center activity relative to each frequency’s baseline dynamics, wavelet power at each frequency was z-scored independently across time within each recording.

ACh oscillation magnitude was quantified for each frequency band (Delta: 0.5–4⍰Hz; Theta: 4–8⍰Hz) in the time–frequency spectra, by computing the signed area under the curve (AUC) of the z-scored power averaged overtime within predefined time windows (before and after treatment) for each mouse and session. Because power was z-scored independently at each frequency, band-limited signed AUC values reflect the net engagement or suppression of that frequency range relative to each animal’s typical activity. These AUC values were used for group-level statistical comparisons across all experimental conditions and treatment history.

*The temporal structure of oscillatory activity* was quantified using burst analysis of the delta and theta bands, where sustained episodes of elevated band-limited activity could be robustly defined. For each recording, z-scored wavelet power was averaged across delta (0.5–4⍰Hz) and theta (4–8⍰Hz) frequencies to generate band-specific power time series. These signals were down-sampled from 50MHz to 10⍰Hz prior to feature extraction. Oscillatory bursts were defined as periods during which band-averaged z-scored power exceeded a fixed threshold of 1 standard deviation above the within-recording mean (zM>M1). To exclude transient fluctuations, only events exceeding a minimum duration were retained (0.5⍰s for delta; 0.25⍰s for theta). From these events, burst fraction (proportion of recording time above threshold) and burst duration (mean length, in seconds, of all suprathreshold epochs) were calculated for each recording. In parallel, the regularity of band-limited oscillatory activity was quantified directly from the fluorescence signal using a rhythmicity index (RI). For each recording, ΔF/F traces were band-pass filtered within the delta and theta ranges using zero-phase digital filtering. The normalized autocorrelation function of the band-passed signal was then computed, and the positive-lag segment corresponding to the expected oscillatory period (delta: 0.25–2⍰s; theta: 0.125–0.25⍰s) was analyzed. RI was defined as the difference between the maximum and minimum autocorrelation values within this interval (trough-to-peak measure), providing an estimate of oscillatory regularity that is independent of signal amplitude. Higher RI values indicate more consistent, periodic oscillatory structure, whereas lower values reflect less regular or more aperiodic activity.

*Event detection and inter-event interval analysis* Discrete ACh transients were identified from ΔF/F photometry signals using a threshold-based peak detection approach. Raw signals were first smoothed using a Gaussian filter (σ = 0.2 s) to reduce high-frequency noise. A slowly varying baseline was estimated using a moving median filter (10 s window) and subtracted to obtain a detrended signal. Noise level was estimated as the median absolute deviation (MAD) of the detrended trace, and events were defined as peaks exceeding a threshold of 2 × MAD. To avoid detecting closely spaced fluctuations as separate events, a minimum peak distance of 0.01 s was imposed. For each detected peak, event onset and offset were defined as the nearest points preceding and following the peak at which the detrended signal crossed zero. Events shorter than 0.02 s were excluded from further analysis. Event duration was calculated as the time between onset and offset, amplitude as the maximum value within the event window, and area as the integral of the detrended signal over the event duration. Inter-event intervals (IEIs) were defined as the time between consecutive event peaks. Event rate was computed as the number of detected events per minute of recording. To quantify temporal variability, the coefficient of variation (CV) of IEIs was calculated for each recording as the ratio of the standard deviation to the mean IEI. All event-based metrics were computed independently for each recording and subsequently used for statistical comparisons across conditions.

### Rating of Dyskinetic Movements

Mice were video-recorded during the imaging sessions to assess their motor behavior. During each photometry session, 3 digital cameras were placed on top, side and bottom of the test cage and mice were video recorded for the duration of the experiment at a video frame rate of 30 frames/s. Videos were analyzed and scored offline. The Abnormal Involuntary Movements (AIMs) is a well-established semi-quantitative scale useful for the assessment of the presence and severity of dyskinesia, as well as its anatomical manifestation ^[18]^. In brief, each animal is assessed for 60 s at 10 min intervals for the whole duration of the session for dyskinetic movements and postures in the Axial (trunk, neck), Limb (hindlimb, forelimb), or Orolingual (face, mouth, tongue) planes. A score of 0-4 is given for each plane based on the following criteria: 0=no AIMs present for 1 min; 1=AIMs present for <30 s; 2=AIMs present for >30 s; 3=AIMs present for 1 min continuously, but interrupted by a tap on the behavioral apparatus; or 4=AIMs present for 1 min continuously and not interrupted by a tap on the behavioral apparatus (**Figure 1E**).

### DA Lesion and Fiber Placement Validation

At the conclusion of experiments, mice were transcardially perfused with 1x PBS followed by 4% paraformaldehyde. To preserve fiber tracts and minimize tissue distortion around the implant site, brains were kept in the skull with fibers in place and post-fixed in 4% paraformaldehyde for 20h before removal. Fixed tissue was sectioned using a freezing microtome (Leica SM 200R, Buffalo Grove, IL, USA). Adjacent sections were processed for immunohistochemical staining for tyrosine hydroxylase (TH) in the substantia nigra pars compacta (SNc) to verify the extent of dopaminergic lesions, or used for direct fluorescence imaging to confirm optical fiber placement and GRAB-ACh sensor expression in the dorsolateral striatum (DLS). All sections were imaged using confocal microscopy with a 10× objective (**Figure 1A-C**).

### Statistical analyses

Statistical analyses were performed using GraphPad Prism version 10.4.1 for Windows (GraphPad Software, Boston, Massachusetts, USA). Differences in total spectral power, band-specific AUC, burst fraction and RI across lesion conditions (DA-lesioned, non-lesioned, sham) were assessed using mixed-effects models with condition as a fixed effect and animal as a random intercept, run separately for each frequency band. Post hoc comparisons were performed using the Holm–Šídák method, restricted to pre-specified contrasts of each control condition against the lesioned hemisphere. Dyskinesia severity across the treatment timeline was compared using a one-way repeated measures ANOVA with Tukey’s multiple comparisons test. Differences in ALO scores between lesioned and sham animals at individual timepoints were assessed using the Mann-Whitney U test. The effects of acute and chronic L-DOPA on ACh oscillatory power and RI were evaluated using mixed-effects models with drug (OFF- vs ON L-DOPA) and condition as fixed effects and animal as a random intercept, run separately for each frequency band. To directly compare the effect of L-DOPA between acute and chronic treatment sessions, a mixed-effects model with session (day 1 vs. day 10) and condition as fixed effects was applied to ON L-DOPA recordings, with post hoc comparisons performed using Šídák’s multiple comparisons test between timepoints within each condition. IEI distributions of ACh events before starting L-DOPA treatment and ON L-DOPA at day 10 of treatment were compared with Kolmogorov-Smirnov test, while Mann Whitney test was used to compare other event metrics. The effect of amantadine on L-DOPA-induced ACh changes was assessed using a two-way ANOVA with amantadine pretreatment and L-DOPA as factors. Within amantadine-pretreated animals, differences between OFF and ON L-DOPA windows were assessed using the Wilcoxon matched-pairs signed rank test. The effect of amantadine alone on baseline ACh oscillations in DA-lesioned hemispheres was assessed using a paired t-test comparing baseline and amantadine windows. For all band-specific analyses, independent models were run for each frequency band without correction across bands, given a priori predictions based on prior work on spontaneous ACh dynamics. A trend toward significance was defined as 0.05 < P ≤ 0.10. Statistical significance was set at P < 0.05.

## RESULTS

### Dopamine depletion shifts striatal ACh dynamics toward increased phasic activity and reduced delta rhythmicity

Injections of 6OHDA in the MFB produced up to 70% loss of nigral dopaminergic cells (**Figure 1A- B**). To determine how loss of dopaminergic input affects striatal intrinsic ACh dynamics, we expressed GRAB-ACh3.0 sensors in the dorsolateral striatum (**Figure 1C**) and recorded spontaneous ACh fluctuations in vivo using bilateral fiber photometry in 6-OHDA–lesioned mice and sham controls (**Figure 2A**). In lesioned animals, signals were acquired from both the DA-depleted and the contralateral intact hemisphere, enabling within-animal comparisons alongside comparisons with sham mice. In intact striatum (sham and non-lesioned hemispheres), spontaneous ACh release was characterized by dominant low-frequency oscillations, with peak power in the 0–2 Hz (delta) range (**Figure 2B**). In contrast, DA-depleted hemispheres exhibited a rightward shift in the power spectral density, with greater relative power distributed across higher frequencies (**Figure 2B**). This reflects a shift in the coordination of spontaneous cholinergic signaling.

**Figure 2.**
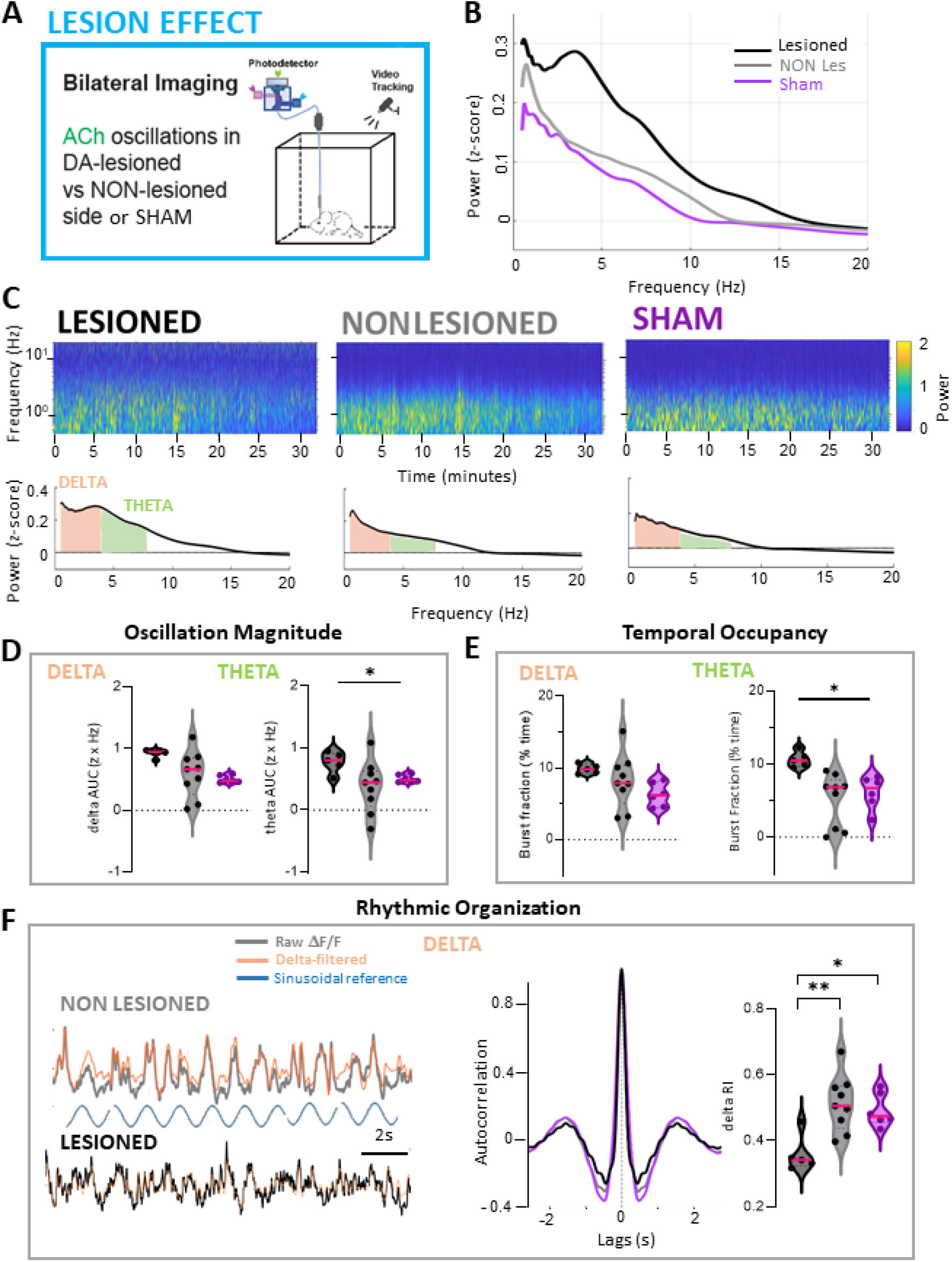
Dopamine depletion alters the spectral profile and reorganizes the temporal structure of striatal ACh dynamics. **(A)** Experimental design. **(B)** Z-scored wavelet power averaged over time is shown in the frequency domain across lesioned, non-lesioned, and sham hemispheres. **(C)** (Bottom) Same z-scored power representation as above with AUC highlighted in the delta (orange) and theta [15] bands. (Top) Time–frequency representations of average power spectral density displayed as colormaps across conditions. **(D–F)** Quantification of band-specific ACh dynamics. Independent mixed-effects models with animal as random intercept were performed for each band (no correction applied across frequency bands). **(D)** AUC analysis revealed a significant effect of condition selectively in the theta band, with increased power in lesioned hemispheres relative to both non-lesioned and sham controls. **(E)** Burst metric analysis showed increased delta burst duration in lesioned hemispheres compared to both control conditions (LES vs. NON LES: P=0.01, LES vs SHAM: P=0.001) and increased theta burst fraction in lesioned hemispheres compared to both control conditions (LES vs. NON LES: P=0.044, LES vs SHAM: P=0.044, Holm-Šídák’s post-hoc multiple comparisons test). **(F)** (Left) Representative ΔF/F traces from non-lesioned and lesioned hemispheres with corresponding delta-band filtered signals overlaid. A sinusoidal reference waveform centered within the delta frequency range is shown for comparison. (Right) Rhythmicity index (RI), computed from the autocorrelation of band-pass filtered ΔF/F signals, revealed reduced delta-band rhythmicity (LES vs. NON LES: P=0.03, LES vs SHAM: P=0.004, Holm-Šídák’s post-hoc multiple comparisons test) in lesioned hemispheres compared to controls. Data are presented as violin plots with individual points representing single hemispheres and median indicated by a horizontal magenta line. The non-lesioned group includes hemispheres from animals with suboptimal lesions that were excluded from subsequent pharmacological analyses. Lesioned n = 9, non-lesioned n = 5, sham n = 6

To better characterize the changes in ACh fluorescence dynamics, we used three complementary analyses to capture: 1) oscillation magnitude in each frequency band, 2) the temporal occupancy of oscillatory activity and 3) the degree of rhythmic organization within each frequency band ^[19].^

To resolve frequency-specific oscillation magnitude, signed area under the curve (AUC) of zscore power spectra was computed separately for two frequency bands partitioned according to standard electrophysiological conventions (delta: 0.5-4Hz and theta: 4-8Hz, **Figure 2C**) and analyzed using independent mixed-effects models per band. A significant effect of condition was detected specifically in the theta band (F (1.214, 5.461) = 6.692, P = 0.04), with post hoc comparisons confirming higher theta power in lesioned hemispheres relative to both control conditions (both P=0.03, Šídák’s multiple comparisons test, **Figure 2D**). No significant effects were detected in the delta band under this analytical framework, indicating that the amplitude of ACh oscillation is enhanced at higher frequencies in lesioned compared to control hemispheres.

To further characterize the temporal organization of ACh oscillatory activity, we used two complementary measures: burst analysis within the delta and theta bands quantified the prevalence of band-limited activity (**Figure 2E**) and an autocorrelation-based rhythmicity index (RI), computed from band-pass filtered ΔF/F signals (**Figure 2F**) which quantifies the degree of periodic temporal structure by measuring how consistently activity recurs at a given timescale, independent of amplitude. Burst analysis revealed that DA-lesioned hemispheres exhibited a significant increase in theta burst fraction (F(1.381, 6.213) = 7.427, P = 0.03) with no significant differences in delta burst fraction (F(1.404, 6.318) = 2.585, P = 0.15, **Figure 2E**) relative to both control conditions, reflecting more dynamically recurring theta activity in lesioned hemispheres. Moreover, in intact striatum delta band-filtered ΔF/F signals exhibited pronounced and regularly spaced autocorrelation peaks, indicative of regular low-frequency oscillatory structure (Figure 2F), which was significantly reduced by DA depletion, as reflected by a decrease in RI (F(1.317, 11.20) = 8.360, P = 0.01, **Figure 2F**). In contrast, theta dynamics were not organized as sustained oscillations (**Supplementary Figure 1**). Together, the reduction in delta rhythmicity, coupled with increased theta power and burst fraction, supports a coordinated reorganization of ACh dynamics toward more frequent, non-oscillatory, higher-frequency activity that disrupts the periodic organization of slow delta dynamics.

### Acute L-DOPA uniformly suppresses power but not rhythmicity of ACh oscillations across conditions

To investigate how L-DOPA exposure influences striatal ACh dynamics, we treated 6-OHDA and SHAM-lesioned mice with low doses of LDOPA (1mg/Kg) for 10 consecutive days and compared the effects of acute (d1, Figure 3) and chronic (d10, Figure 4) L-DOPA administration.

**Figure 3.**
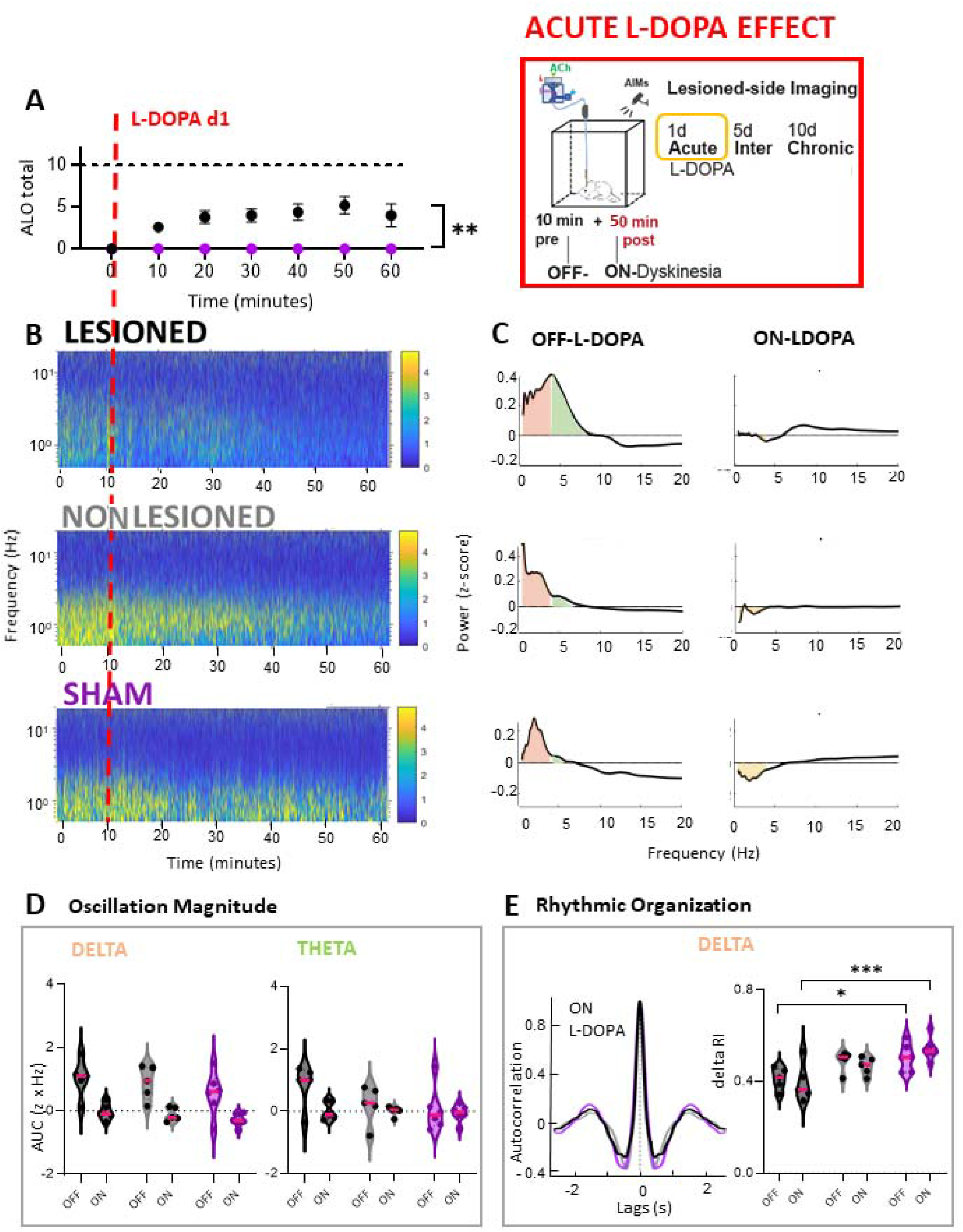
Acute L-DOPA suppresses the power, but not the rhythmicity, of ACh oscillations regardless of DA depletion status. **(A)** Dyskinesia scores are expressed as total ALO over time following acute L-DOPA injection. Lesioned animals (black) showed significantly higher ALO scores than sham controls (purple, P=0.005, Mann-Whitney test). **(B)** Average striatal ACh power spectral density is shown as colormaps and **(C)** Z-scored power averaged over baseline time window (OFF L- DOPA: first 10 minutes) and over time window with stable dyskinesia (ON L-DOPA: 30-50 min from recording onset) is shown in frequency domain with AUC highlighted in the delta band. **(D)** Acute L- DOPA significantly reduced oscillatory power in the delta and theta bands (main effect of drug: delta P<0.0001, theta P=0.04), with no significant differences between conditions in any band (ns, drug x condition interactions: delta P=0.85, theta P=0.24; mixed effects model). **(E)** Rhythmicity index (RI), computed from the autocorrelation of band-pass filtered ΔF/F signals, revealed that acute L-DOPA does not rescue the reduced delta-band rhythmicity in lesioned hemispheres, which is significantly lower before L-DOPA (OFF L-DOPA: LES vs. NON LES: P=0.07, LES vs SHAM: P=0.02) and after L-DOPA compared to sham hemispheres (ON-LDOPA: LES vs. NON LES: P=0.07, LES vs SHAM: P=0.02, Šídák’s post-hoc multiple comparisons test). Data are presented as violin plots with individual points representing single hemispheres and median indicated by a horizontal magenta line. Lesioned and non lesioned n=5, sham n=6

**Figure 4.**
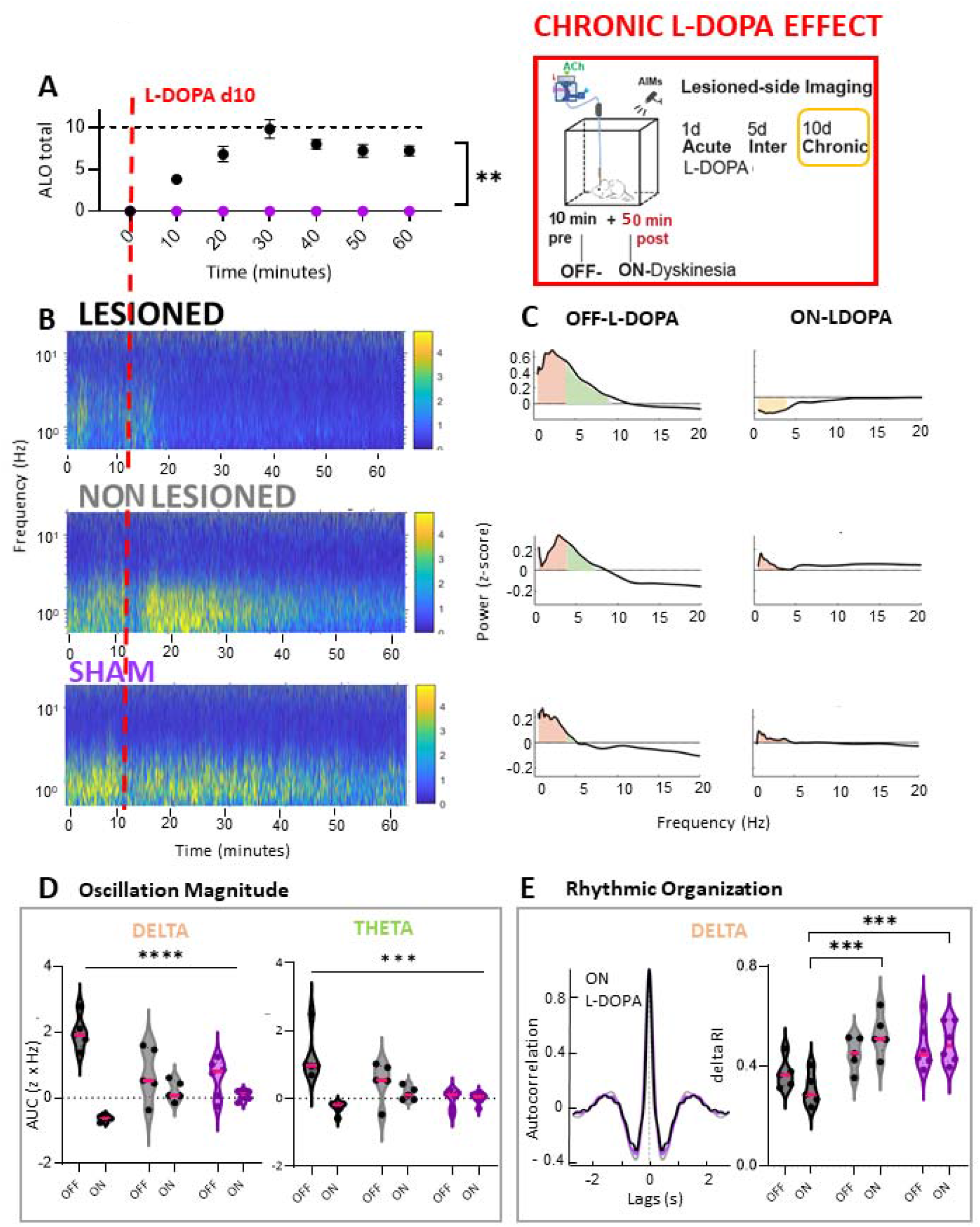
Chronic L-DOPA induces lesion-dependent disruption of striatal delta ACh oscillations. **(A)** Dyskinesia scores are expressed as total ALO over time following acute L-DOPA injection. Lesioned animals (black) showed significantly higher ALO scores than sham controls (purple, P=0.004, Mann- Whitney test). **(B)** Average striatal ACh power spectral density is shown as colormaps and **(C)** Z- scored power averaged over baseline time window (OFF L-DOPA: first 10 minutes) and ON LDOPA (30-50 min from recording onset) is shown in frequency domain with AUC highlighted in the delta band. **(D)** Chronic L-DOPA, associated with established dyskinesia, revealed significant disruption of oscillatory activity across all frequency bands (main effect of drug: delta P<0.0001, theta P=0.0003), with significant differences across lesion status (drug × condition interactions: delta p<0.0001, theta p=0.0006, mixed effects model). **(E)** Rhythmicity index (RI), computed from the autocorrelation of band-pass filtered ΔF/F signals, revealed that chronic L-DOPA preferentially depressed delta-band RI in lesioned hemispheres compared to controls (ON-LDOPA: LES vs. NON LES: P=0.0002, LES vs SHAM: P=0.01. Data are presented as violin plots with individual points representing single hemispheres and median indicated by a horizontal magenta line. Same animals as above

Acute L-DOPA treatment elicited significantly higher contralateral rotations and abnormal involuntary movements, scored in the axial, orolingual and limb (ALO) dimension, in lesioned animals compared to sham controls (P=0.005, Mann-Whitney test, **Figure 3A**), confirming that abnormal involuntary movements were specific to DA depletion. The slow onset and mild severity of the AIMs in the acute d1 phase are consistent with an initial sensitization.

In the striatum, acute L-DOPA produced a robust suppression of ACh oscillatory power in the delta and theta bands (main effect of drug: delta F (1, 26) = 33.43, P<0.0001, theta F (1, 26) = 4.436, P=0.04, mixed effects model, **Figure 3B–D**). Critically, this suppression was uniform across all conditions, with no drug × condition interactions in any band (delta: F (2, 26) = 0.1647, P=0.85, theta: F (2, 26) = 1.495, P=0.24), indicating that acute dopaminergic restoration suppresses ACh oscillations independently of prior DA depletion status. This effect was specific to L-DOPA, as saline injections did not produce comparable reductions under any condition (**Supplementary Figure 2**).

However, despite the uniform suppression of oscillatory power across conditions, analysis of temporal structure revealed condition- and band-specific effects on the organization of ACh dynamics. Mixed-effects analysis revealed a significant main effect of condition on delta-band rhythmicity (F (2, 13) = 9.923, P=0.002, **Figure 3E**), with lesioned hemispheres exhibiting significantly reduced RI compared to sham both before and after L-DOPA administration (Šídák’s post-hoc multiple comparisons test). This indicates a persistent impairment of delta oscillatory organization in the DA-depleted striatum, which is not re-established by acute L-DOPA. Notably, although acute L-DOPA reduced delta-band power across all conditions, control hemispheres did not exhibit a comparable reduction in rhythmicity, indicating that suppression of oscillatory amplitude alone was not sufficient to disrupt temporal organization.

Together, these findings indicate that acute L-DOPA uniformly suppresses the magnitude of low-frequency ACh oscillations across conditions. Despite this global reduction in oscillatory power, the temporal organization of delta activity is preserved in control conditions but is not restored in the DA lesioned striatum, where it remains impaired following L-DOPA exposure.

### Chronic L-DOPA induces lesion-specific disruption of ACh delta rhythmicity while enhancing ACh activity

At day 10 of L-DOPA treatment, chronically treated DA-lesioned animals displayed enhanced contralateral rotations and well-established dyskinetic behaviors, characterized by sustained dyskinetic postures with ALO scores significantly higher than sham controls (Mann-Whitney test, P=0.004, **Figure 4A**), confirming full expression of LID following repeated treatment. With repeated L-DOPA exposure, dyskinetic behaviors within lesioned animals emerged with earlier onset and reached increased severity compared to acute L-DOPA. ALO scores increased significantly across the treatment timeline (F (1.598, 9.587) = 11.80, P=0.0036, one-way repeated measures ANOVA), with day 1 scores significantly lower than both day 5 and day 10 (both P=0.02, Tukey’s multiple comparisons), while day 5 and day 10 did not differ significantly (P=0.4), indicating that dyskinesia severity plateaued by mid-treatment.

Strikingly, chronic L-DOPA in animals with established dyskinesia produced a pattern of ACh disruption that differed from the uniform effects observed with acute treatment. Although L-DOPA continued to suppress low-frequency oscillatory power (delta: F(1, 26) = 52.34, P < 0.0001; theta: F(1, 26) = 17.95, P = 0.0003; **Figure 4B–D**), this effect was no longer uniform across conditions, as indicated by significant drug × condition interactions in both bands (delta: F(2, 26) = 16.40, P < 0.0001; theta: F(2, 26) = 9.998, P = 0.0006; **Figure 4B–D**). These findings indicate that, with repeated treatment, the impact of L-DOPA on ACh oscillation magnitude becomes dependent on prior DA depletion. Consistent with this divergence, delta-band temporal organization, quantified by the RI, also exhibited a significant drug × condition interaction (F(2, 13) = 6.791, P = 0.01), with post hoc comparisons revealing that lesioned hemispheres had lower delta RI than both control conditions after chronic L-DOPA administration (**Figure 4E**).

To determine whether the effects of L-DOPA on ACh dynamics evolved across repeated treatment in a lesion-dependent manner, we directly compared ON-L-DOPA frequency band power and RI between day 1 and day 10 using a mixed-effects model with session and condition as fixed effects (**Figure 5**). In the delta band, a significant session × condition interaction was detected (F(2, 26) = 16.98, P < 0.0001, **Figure 5A**), revealing opposing trajectories across groups. Specifically, lesioned hemispheres exhibited a significant decrease in delta power from day 1 to day 10 (P = 0.0004, Šídák’s multiple comparisons test), indicating a progressive enhancement of ACh suppression with chronic L-DOPA. In contrast, sham hemispheres showed a significant increase in delta power across sessions (P = 0.02), consistent with attenuation of the acute suppressive effect in the absence of DA depletion, while non-lesioned hemispheres showed a similar trend (P = 0.08). Theta-band power showed similar opposing trajectories but only a trend toward a significant session × condition interaction (F(2, 26) = 2.857, P = 0.08, **Figure 5B**).

**Figure 5.**
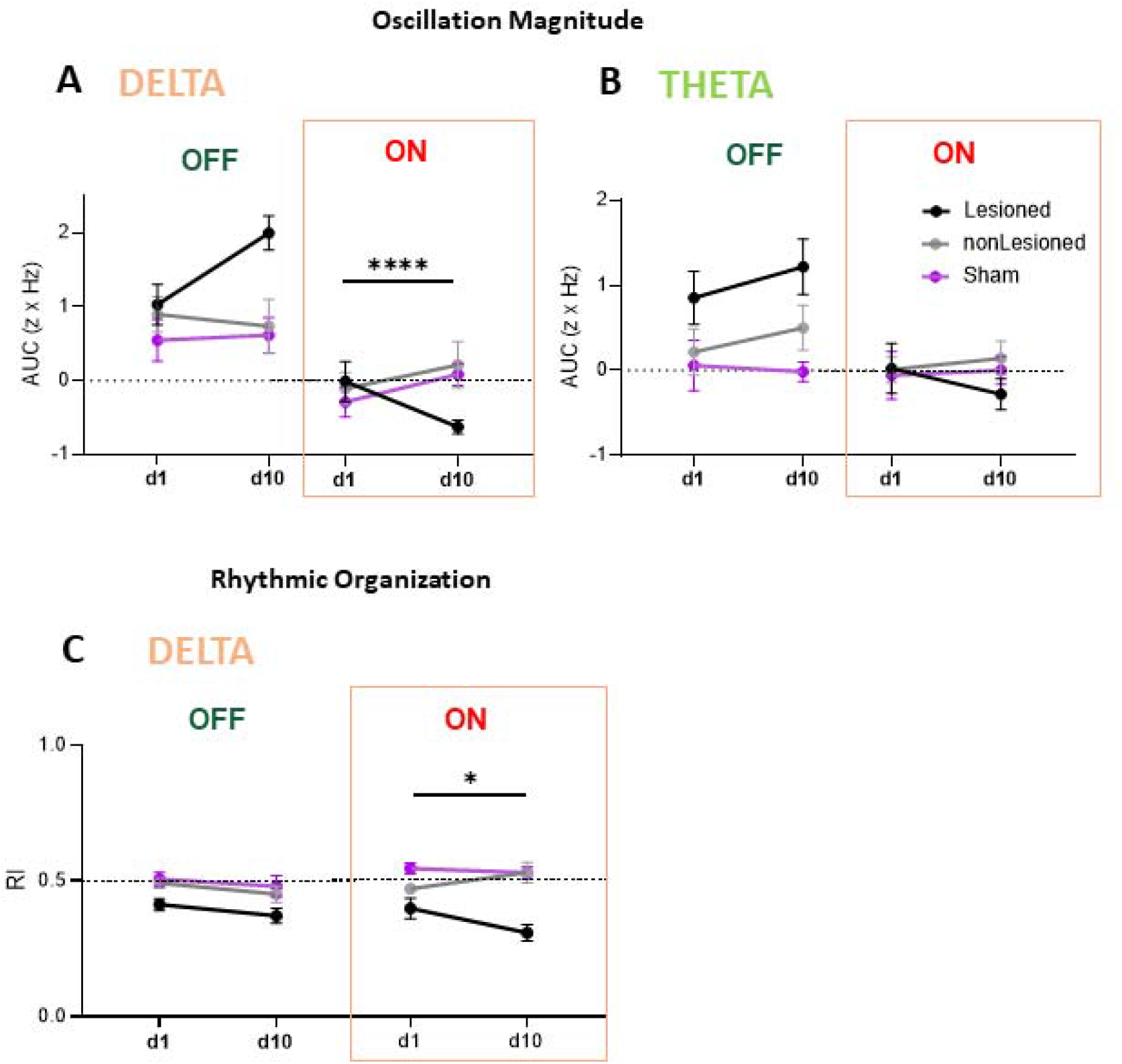
Direct comparison between acute and chronic L-DOPA treatment on striatal ACh oscillations across lesion conditions. **(A)** Delta and **(B)** theta-band AUC levels OFF and ON-LDOPA at day 1 and day 10, shown for lesioned (black), non-lesioned [30], and sham (purple) hemispheres. Lines connect individual session means within each condition. **(C)** Same comparisons for delta RI

A similar pattern was observed for delta-band RI, which also exhibited a significant session × condition interaction (F (2, 13) = 4.787, P=0.03) with lesioned and control hemispheres showing opposing longitudinal changes that paralleled those observed for power (**Figure 5C**). These results demonstrate that chronic L-DOPA exposure progressively disrupts the temporal organization of striatal ACh dynamics selectively in the DA-depleted hemisphere in association with dyskinesia manifestation. Importantly, both delta-band oscillatory activity and theta-band bursts returned during OFF L-DOPA recordings throughout the treatment timeline. This is supported by the absence of significant session × condition interactions in AUC for both delta (F(4, 26) = 2.221, P = 0.09) and theta bands (F(4, 26) = 1.236, P = 0.3; **Figure 5A-B**), as well as in delta-band RI (F(4, 26) = 0.8462, P = 0.5; **Figure 5C**). Notably, lesion-associated dysfunctions of ACh dynamics, including increased theta activity (condition effect: F (2, 17) = 8.402, P=0.003) and reduced delta RI (F (2, 17) = 9.901, P=0.001) were also preserved during OFF L-DOPA recordings even after chronic treatment (**Figure 5B-C**). Together, these findings indicate that the collapse of delta oscillatory structure reflects an effect of DA replacement that is state-dependent and associated with LID, while the changes in intrinsic cholinergic network function induced by DA deprivation are permanent and not reversed by DA replacement.

While spectral and rhythmicity analyses revealed marked disruptions in the temporal organization of ACh signaling, these measures do not directly inform whether the overall occurrence of ACh transients is increased or decreased. Given the prevailing view that cholinergic activity is elevated following L-DOPA treatment, we next quantified the occurrence of discrete ACh events independently of oscillatory structure. Event-based analysis revealed that chronic L-DOPA treatment compared to before L-DOPA exposure in lesioned hemispheres shifted the distribution of inter-event intervals (IEIs) toward shorter values (P=0.0002, Kolmogorov-Smirnov test, **Figure 6 A-B**), indicating more frequent event occurrence, consistent with increased rate of ACh transients (P=0.03, Mann Whitney test, **Figure 6B**). In addition, the coefficient of variation of IEIs (CV_IEI) was significantly increased (P=0.015, Mann Whitney test, **Figure 6C**), reflecting greater temporal irregularity in event timing. In contrast, these effects were not observed in sham hemispheres (**Figure 6D**), where L-DOPA did not significantly alter IEI distribution (P=0.13), event rate (P=0.09, **Figure 6E**) and variability (P=0.9, **Figure 6F**), indicating that the impact of L-DOPA on event frequency depends on the DA-depleted state.

**Figure 6.**
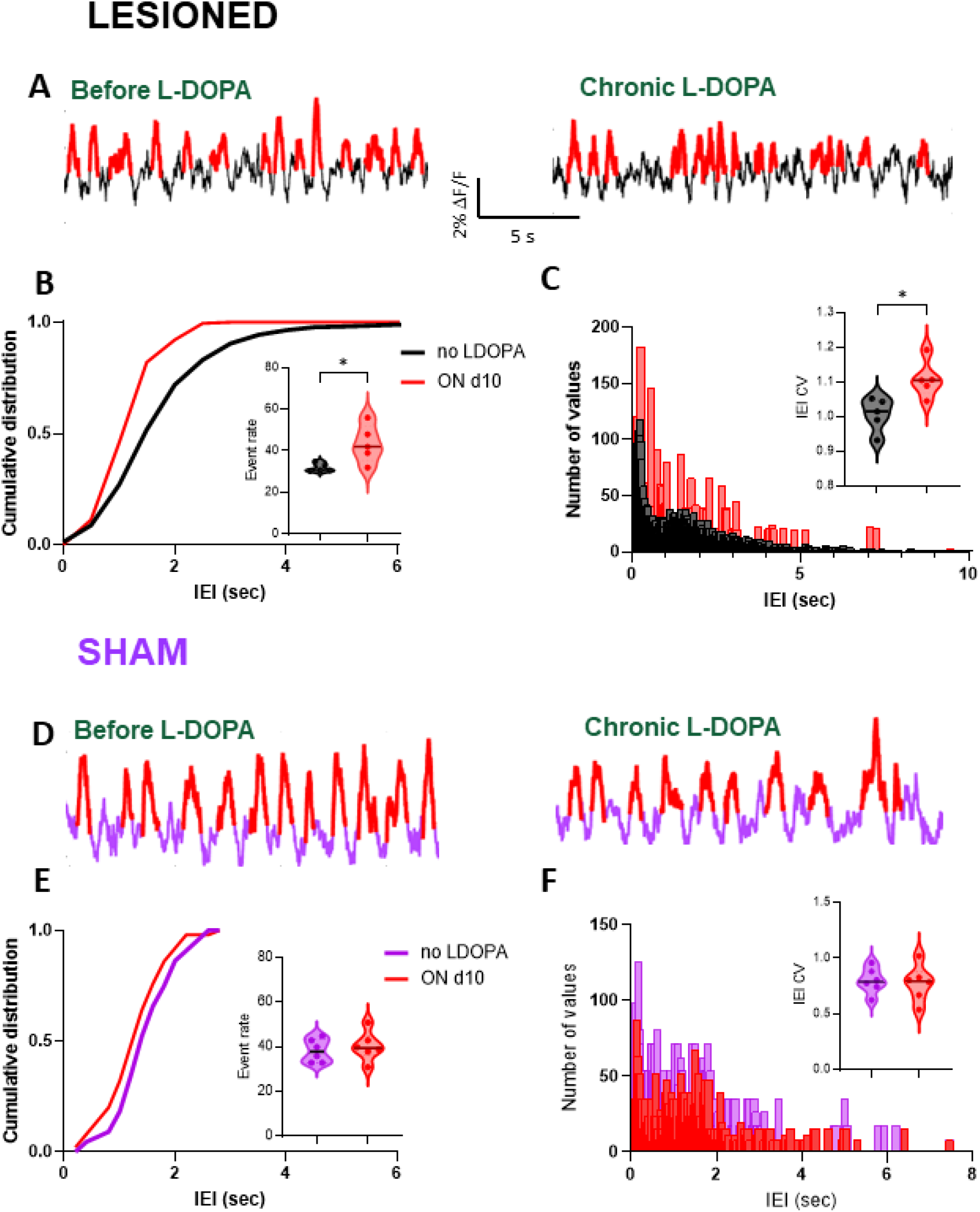
Chronic L-DOPA increases frequency and temporal irregularity of ACh transients in the lesioned striatum. **(A)** Representative ΔF/F traces from lesioned hemispheres before and after chronic L-DOPA treatment, with detected events highlighted in red. **(B)** Distribution of inter-event intervals (IEI) shows a leftward shift following chronic L-DOPA (red), indicating shorter intervals between events and increased event rate (insert). **(C)** IEI histogram and elevated coefficient of variation (CV, insert) following chronic L-DOPA, reflect increased temporal irregularity of ACh transients. **(D-F)** Same event-based analysis in sham hemispheres before and after chronic L-DOPA treatment. Data are presented as violin plots with individual points representing single hemispheres and median indicated by a horizontal line. Statistical comparisons were performed using Kolmogorov-Smirnov test and Mann Whitney test. n =5, lesion; n=6, sham hemispheres

Together, these findings indicate that L-DOPA increases the occurrence of ACh transients while simultaneously degrading their temporal coordination.

### Amantadine attenuates dyskinesia and preserves low-frequency striatal ACh oscillations during L- DOPA treatment

To determine whether attenuation of dyskinesia expression would influence striatal Ach dynamics during L-DOPA treatment, we examined the effects of the anti-dyskinetic drug amantadine in chronically L-DOPA–treated mice (**Figure 7**). Consistent with its established clinical profile, amantadine (60 mg/kg, administered 100 minutes before L-DOPA)^[20]^ significantly reduced the severity and duration of abnormal involuntary movements (Mann-Whitney test, P=0.02, **Figure 7A**). At the ACh network level, amantadine prevented the disruption of low-frequency ACh oscillations observed during dyskinetic states under chronic L-DOPA alone (**Figure 7B-C**). Within amantadine-pretreated animals, no significant change in delta or theta power was detected between OFF and ON L-DOPA windows (P=0.5 and P=0.75 respectively, Wilcoxon matched-pairs signed rank test), indicating that amantadine abolished L-DOPA-induced suppression of ACh oscillations observed with chronic treatment. Direct comparison with chronic L-DOPA alone confirmed a significant amantadine × L-DOPA interaction in the delta band (F (1, 6) = 15.28, P=0.008, two-way repeated measures ANOVA), with a significant main effect of L-DOPA (F (1, 6) = 35.50, P=0.001) and no independent effect of amantadine (F (1, 6) = 0.003035, P=0.95, **Figure 7D**). A trend toward interaction was observed in the theta band (F (1, 6) = 4.738, P=0.07), likely reflecting limited power given the sample size (**Figure 7E**).

**Figure 7.**
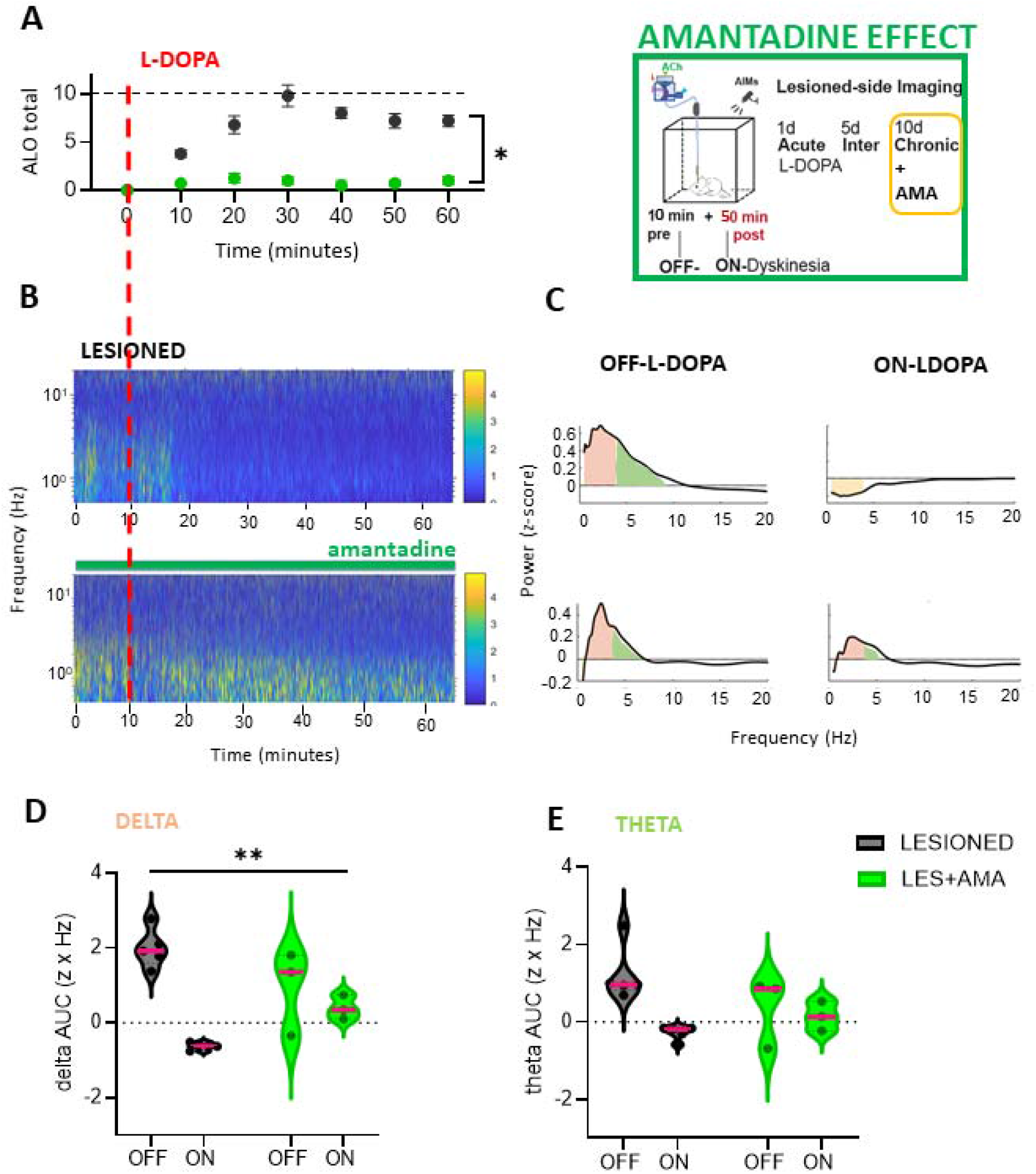
Amantadine preserves low-frequency ACh oscillations during L-DOPA treatment. **(A)** ALO dyskinesia scores in amantadine-pretreated [15] and lesioned animals chronically treated with L-DOPA (black, same as Figure 4A). Pre-treatment with amantadine lowered ALO scored significantly compared with L-DOPA alone (P=0.02, Mann-Whitney test). **(B)** Colormaps of average striatal ACh power spectral density from lesioned hemispheres, as in Figure 4B, and when pre-treated with amantadine. **(C)** Z-scored power averaged over OFF L-DOPA (first 10 min) and ON L-DOPA (30–50 min from recording onset) windows, with AUC highlighted in the delta and theta bands, showing preserved power in both frequency bands ON L-DOPA in the presence of amantadine. **(D)** Direct comparison with chronic L-DOPA alone confirmed a significant amantadine × L-DOPA interaction in the delta band and **(E)** a similar trend in the theta band

Notably, amantadine alone was associated with a reduction of theta ACh oscillations (P=0.02, paired t-test, **Supplementary Figure 3**) that are enhanced during OFF L-DOPA periods in DA-lesioned hemispheres (see also Figure 2), suggesting that glutamatergic modulation may influence the oscillatory organization of striatal ACh signaling both before and during L-DOPA treatment. Together, these findings indicate that the anti-dyskinetic efficacy of amantadine is accompanied by preservation of ACh temporal dynamics in the DA-depleted striatum and motivate further investigation into how glutamatergic and network-level modulation shapes the oscillatory organization of striatal cholinergic signaling.

## DISCUSSION

This study examined how DA depletion and L-DOPA treatment reshape the oscillatory organization of striatal ACh signaling in awake behaving mice. Rather than focusing on mean transmitter levels or single-cell intrinsic properties, we characterized state-dependent changes in the spectral structure of endogenous ACh dynamics across PD- and LID-relevant conditions. We found that DA depletion and L-DOPA treatment differentially reshape the spectral organization of striatal ACh signaling. DA loss shifted ACh dynamics toward higher-frequency, non-oscillatory activity and reduced regularity of low frequency oscillations (**Figure 2**), acute L-DOPA broadly suppressed low-frequency rhythms independent of lesion status and abolished lesioned-enhanced theta frequency fluctuations (**Figure 3**), chronic L-DOPA disrupted low-frequency synchronization selectively in the DA-depleted striatum during dyskinesia (**Figure 4-5**), while elevating ACh transient frequency and variability (**Figure 6**). Finally, amantadine preserved low-frequency oscillatory structure while reducing LID (**Figure 7**).

### Dopamine Depletion Effects

Both DA loss and L-DOPA treatment have been associated with increased cholinergic tone and enhanced cholinergic interneuron (ChI) excitability ^[21–24]^. Our findings extend this framework by showing that DA depletion does not simply increase cholinergic activity, but reshapes its temporal organization. Specifically, loss of dopaminergic input shifts ACh dynamics toward faster timescales, increasing activity in the theta frequency-band while weakening the regularity of slow delta oscillations (**Figure 2**). This pattern suggests a reorganization of cholinergic population dynamics, in which activity becomes more frequently and consistently engaged at higher frequencies, while slow fluctuations lose temporal stability.

These observations introduce a network-level dimension to cholinergic dysregulation in PD-like conditions that extends beyond changes in single-cell excitability or average transmitter levels. Nonetheless, the increased phasic activity and reduced delta oscillatory organization observed here are broadly consistent with prior experimental and computational studies reporting elevated firing rates and reduced regularity in ChIs following DA depletion ^[25] [26–29]^. In this context, our spectral analyses provide a population-level readout that is sensitive to changes in underlying cellular dynamics, although they do not directly measure neuronal firing or synaptic mechanisms.

From a functional perspective, a substantial body of experimental and clinical work has linked increased striatal cholinergic signaling to PD symptomatology. While the present approach quantifies relative spectral features rather than absolute ACh levels, the observed increase in high-frequency, phasic activity is consistent with periods of enhanced cholinergic signaling in the DA-depleted state. Notably, this non-oscillatory activity is sensitive to L-DOPA treatment (**Figure 3-4**) yet persists during OFF L-DOPA recordings even after chronic exposure (**Figure 5B**), indicating that this fast cholinergic component remains a stable feature of the DA-depleted state while retaining sensitivity to dopaminergic tone. This pattern raises the possibility that L-DOPA–dependent depression of this activity contributes to its therapeutic effects on motor function, whereas its re-emergence during wearing-off periods may participate in reappearance of PD symptoms.

Future studies using lifetime-based photometry, which enables more direct estimation of baseline neurotransmitter concentration, will be important for determining how changes in oscillatory structure relate to absolute ACh levels.

### Acute L-DOPA Effects

Acute L-DOPA produced a broad suppression of low-frequency ACh oscillation magnitude across all conditions, independent of lesion status, while preserving the intrinsic temporal organization of delta activity in DA intact hemispheres (**Figure 3**). This is consistent with the known ability of dopaminergic signaling to inhibit ChI activity and ACh release through both direct mechanisms (e.g., activation of inhibitory D2 receptors on ChIs) and indirect circuit effects, such as recruitment of GABAergic interneurons. These findings therefore point to a strong pharmacological suppression of ACh output rather than a global reorganization of cholinergic dynamics ^[28, 30]^. Notably, acute L-DOPA reduced oscillatory magnitude across frequencies, normalizing the lesion-induced enhanced theta components without restoring delta-band disorganization. This dissociation indicates that DA replacement acutely suppresses ACh activity but it is insufficient to recover the network-level synchronization underlying slow oscillations.

Therefore, dopaminergic signaling can rapidly modulate the strength of cholinergic output, which may be beneficial to restore normal movement in PD conditions by limiting excessive cholinergic signaling, but restoration of coordinated slow network dynamics requires longer-term circuit reorganization.

### Chronic L-DOPA Effects

In contrast to the uniform suppression observed with acute treatment, chronic L-DOPA exposure, coincident with dyskinesia expression, produced lesion-specific disruptions of ACh dynamics. In the DA-depleted striatum, chronic treatment resulted in a pronounced suppression of both the power and rhythmicity of delta-band ACh oscillations, whereas non-lesioned and sham hemispheres exhibited a comparatively attenuated drug effect (**Figure 4**). These opposing trajectories indicate that repeated L-DOPA exposure fundamentally alters the relationship between dopaminergic signaling and cholinergic dynamics in a lesion-dependent manner.

At the network level, these changes suggest a transition from a synchronized, rhythmically organized cholinergic state to a temporally disorganized regime in the lesioned striatum. Such a collapse of low-frequency oscillatory structure could arise from multiple, non-mutually exclusive mechanisms, consistent with reports of dysregulated and maladaptive cholinergic activity in LID, including altered intrinsic excitability of ChIs ^[31]^, disruption of coordinated pause–rebound dynamics ^[32]^ and aberrant responses to dopaminergic receptor activation ^[33, 34]^. In this context, the reduction in oscillatory activity does not necessarily imply reduced cholinergic activity per se, but rather a loss of coordinated population dynamics. In fact, the observed increase in ACh event frequency combined with greater temporal variability is consistent with excessive but poorly organized cholinergic activity (**Figure 6**). This disorganized activity may impair the ability of the striatum to coordinate motor output, contributing to the emergence of involuntary movements.

Importantly, these effects appear to be again confined to the ON state, with both oscillation power and rhythmicity increasing during the OFF state (**Figure 5**). This suggests that the disorganization of ACh dynamics reflects an acute and maladaptive drug effect rather than a permanent loss of oscillatory capacity. In contrast, the attenuation of ACh suppression observed in non-lesioned and sham hemispheres with chronic L-DOPA treatment is more consistent with homeostatic adaptation or receptor desensitization, which, in response to repeated dopaminergic stimulation, progressively reduce ChI responsiveness through a mechanism potentially involving D2Rs ^[35]^. It is possible that these signaling mechanisms are altered following prolonged L-DOPA exposure in chronically DA depleted states. Interestingly, D2R signaling in ChIs has been shown to paradoxically enhance cholinergic activity in the dorsolateral striatum during the L-DOPA ON state ^[34]^, highlighting dysfunctional D2R responses as a candidate mechanism linking cholinergic dysfunction to dyskinesia pathophysiology ^[36]^.

### Amantadine Effects

Amantadine acts in part as a weak NMDA receptor antagonist, reducing excitatory drive onto striatal neurons, including ChIs. In particular, both thalamic and cortical glutamatergic inputs strongly regulate ChI activity ^[37]^ and excitatory inputs have been implicated in shaping ACh oscillatory patterns ^[9]^. From a circuit-level perspective, the normalization of low-frequency ACh oscillations by amantadine, observed both before and after L-DOPA treatment and in parallel with its anti-dyskinetic effects, suggests a role for glutamatergic inputs in stabilizing cholinergic temporal dynamics. The suppression of theta components with amantadine alone in DA lesioned hemispheres (**Suppl. Figure 3**), suggests that this deficit reflects a circuit-level alteration, consistent with cortico- and thalamo-striatal synaptic adaptations induced by DA loss.

Furthermore, the ability of amantadine to preserve low-frequency ACh oscillations during L-DOPA treatment is consistent with a model in which repeated DA exposure drives excessive or dysregulated excitatory input that contribute to the destabilization of cholinergic network dynamics following DA depletion ^[37]^ (**Figure 7**). Notably, when amantadine prevented this ON-state disruption, dyskinetic behaviors were concurrently reduced, linking preservation of low-frequency ACh organization to improved motor outcomes.

### Potential functional role of delta cholinergic rhythmicity in motor control

A major finding of the present study is that dyskinetic states were associated not simply with altered ACh signaling, but with a profound disruption of its temporal organization. This observation raises the question of whether low-frequency cholinergic oscillations serve a physiological function in the healthy striatum and whether their loss contributes to abnormal motor output.

One possibility is that low-frequency cholinergic rhythms contribute to the basal ganglia’s ability to suppress competing motor programs while facilitating appropriate actions. ChIs are among the most influential modulators of striatal activity because ACh regulates SPNs, interneurons, DA release, and corticostriatal transmission simultaneously. Under normal conditions, slow synchronized cholinergic oscillations may impose recurring windows of inhibition and disinhibition across the striatum. This temporal structure could help ensure that only the most strongly represented motor programs gain control of downstream basal ganglia circuits. In this framework, low-frequency ACh oscillations would stabilize motor selection by constraining the simultaneous activation of competing striatal ensembles. When delta rhythmicity is lost, disruption of this temporal organization may result in weaken motor filtering, allowing inappropriate motor fragments to emerge as involuntary movements.

An alternative, though not mutually exclusive, interpretation is that low-frequency cholinergic rhythms contribute to the temporal organization of automated motor behaviors. For instance, the frequency range of spontaneous cholinergic oscillations closely overlaps with the rhythmic structure of locomotion. Previous studies have demonstrated coupling between delta-frequency membrane potential oscillations in striatal ChIs and the stepping cycle ^[10]^, suggesting that cholinergic rhythmicity may play a role in organizing movement patterning. ChIs receive convergent input from cortical, thalamic, dopaminergic, and local striatal sources. This connectivity places them in an ideal position to integrate information across multiple motor- and non-motor-related pathways and transform it into a temporally structured signal capable of influencing large populations of striatal neurons. Within this framework, low-frequency cholinergic oscillations could represent a network-level signal that helps maintain the timing and coordination of sequential motor commands required for smooth, automated behaviors. Such a mechanism is particularly intriguing in light of the progressive loss of delta-frequency organization observed during dyskinesia, raising the possibility that abnormal involuntary movements reflect not only impaired action selection but also a breakdown in the temporal coordination of motor execution.

Although the present study does not establish a causal role for cholinergic oscillations in shaping the structure of ongoing movement, in action selection or dyskinesia generation, the preservation of low-frequency structure by amantadine together with its anti-dyskinetic effects is consistent with the hypothesis that restoration of cholinergic temporal organization contributes to therapeutic benefit. Future studies combining direct manipulation of cholinergic rhythms with circuit-level recordings and detailed kinematic analyses will be required to determine whether low-frequency ACh oscillations actively regulate motor execution/selection and whether their restoration can prevent the emergence of dyskinetic behaviors.

## CONCLUSIONS

Collectively, our findings reveal that DA depletion and L-DOPA treatment do not simply alter the magnitude of cholinergic signaling, but fundamentally reshape its temporal organization. Across disease and treatment states, striatal ACh dynamics shift from coordinated low-frequency oscillations toward more irregular, higher frequency activity with DA lesion, to a further state-dependent breakdown of network coordination associated with LID. These results suggest that loss of temporal structure is a key feature of striatal dysfunction in Parkinsonian and dyskinetic conditions. We propose that low-frequency cholinergic oscillations represent a network-level signature of coordinated striatal processing required for normal motor selection. Their disruption may reflect a loss of temporal organization within basal ganglia circuits that contributes to dyskinesia. Defining the circuit mechanisms that generate and maintain coordinated cholinergic dynamics, particularly the role of cortico-thalamostriatal inputs, will be essential for understanding how neuromodulatory signals shape motor control in health and disease.

## AKNOWLEDGEMENT

We thank Josh Randolph for his assistance in maintaining and caring for the mouse colony, Dr. David G. Standaert for his constructive feedback on the manuscript and the American Parkinson Disease Association (APDA) for funding this project.

## Author contributions

**SCG**: Investigation, Formal Analysis. **KEJ**: Investigation, Formal Analysis, Writing- Reviewing and Editing. **MS**: Conceptualization, Investigation, Formal Analysis, Data Curation and Visualization, Writing- Original draft preparation, Funding Acquisition, Supervision. All authors read and approved the final manuscript.

## Declarations of interest

The authors have no relevant financial or non-financial interests to disclose. This study was funded by the American Parkinson Disease Association.

---

**Supplementary Figure 1.**
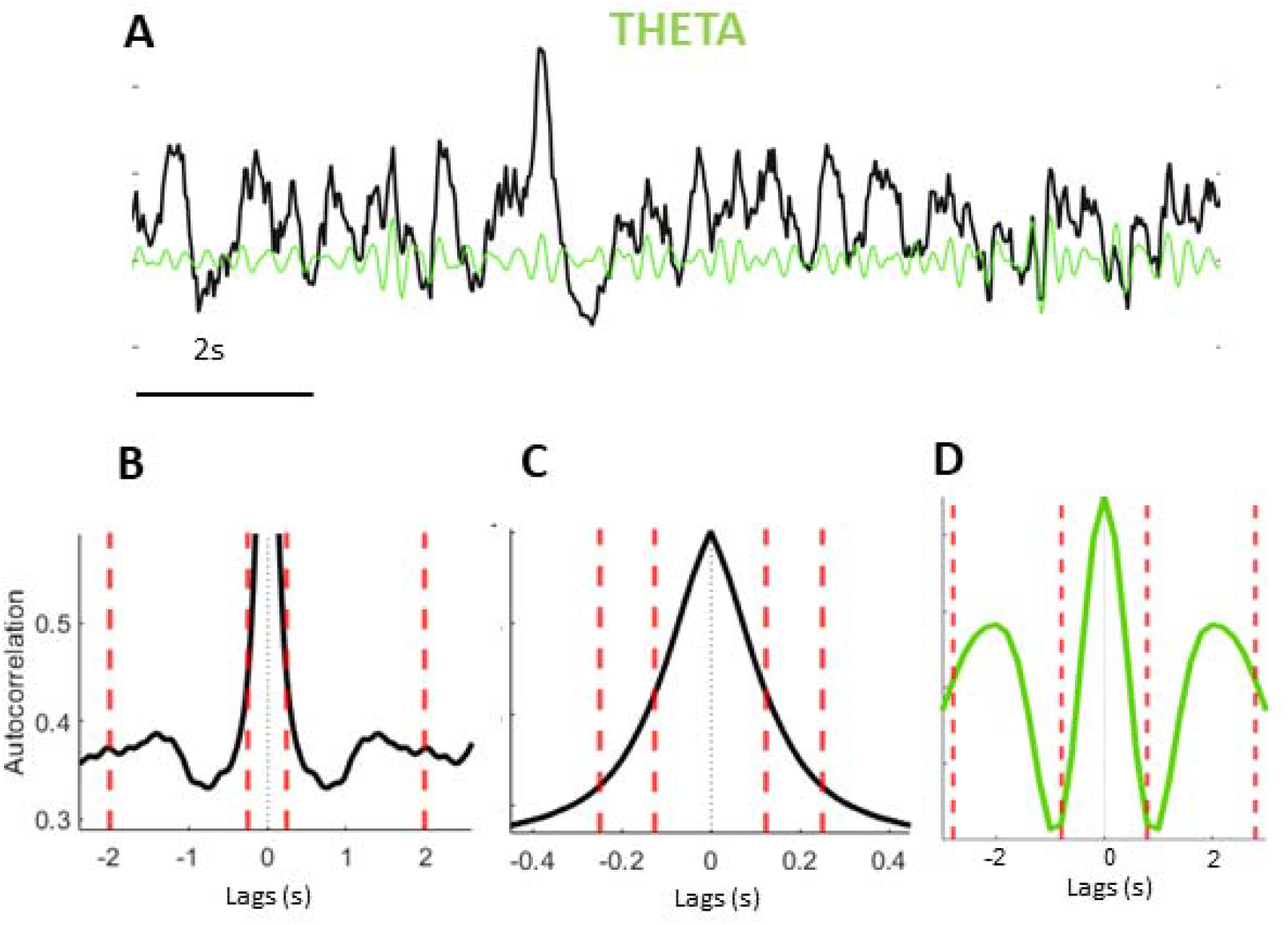
Theta-band activity lacks stable oscillatory structure. **(A)** Representative ΔF/F trace with overlaid theta-band filtered signal (light green), which produces smooth, narrowband fluctuations that are not evident in the raw trace. **(B)** Representative autocorrelations of raw ΔF/F signals for delta and **(C)** theta lag ranges (red dotted lines), illustrating the presence of periodic structure in delta but not theta. **(D)** Representative autocorrelations of theta-filtered ΔF/F showing artificially perfect periodical structure

**Supplementary Figure 2.**
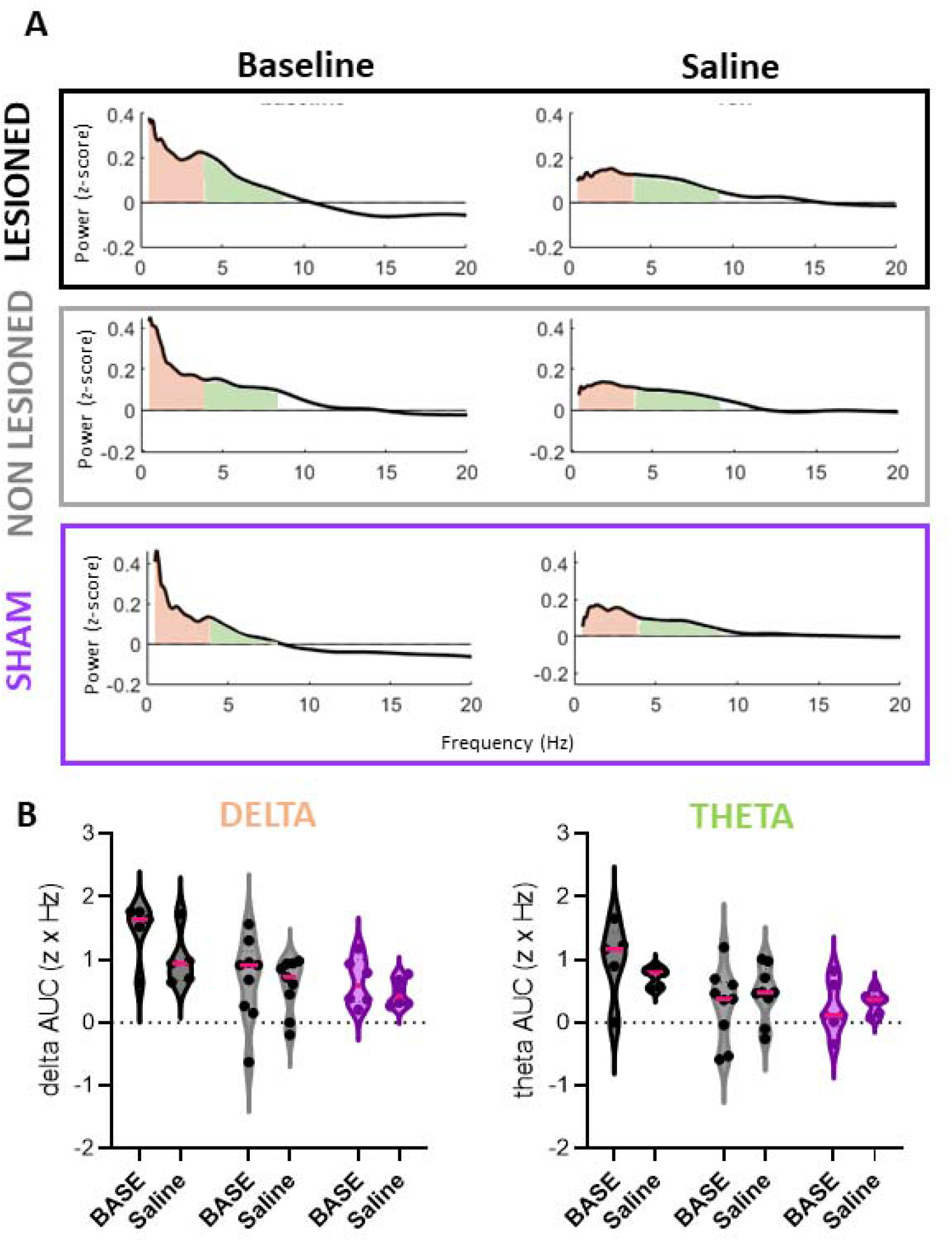
Saline injections do not reduce low frequency ACh oscillations. **(A)** Z-scored power averaged over baseline time window (Baseline: first 10 minutes) and after saline injections (30-50 min from recording onset) is shown in frequency domain with AUC highlighted in the delta band. **(B)** Saline did not produce any change in low frequency oscillations of ACh signal (drug effect: delta F (1, 17) = 2.639, P=0.12, theta F (1, 17) = 0.0008396, P=0.98, Lesioned n=5, non lesioned n=9, Sham n=6, Mixed effects model)

**Supplementary Figure 3.**
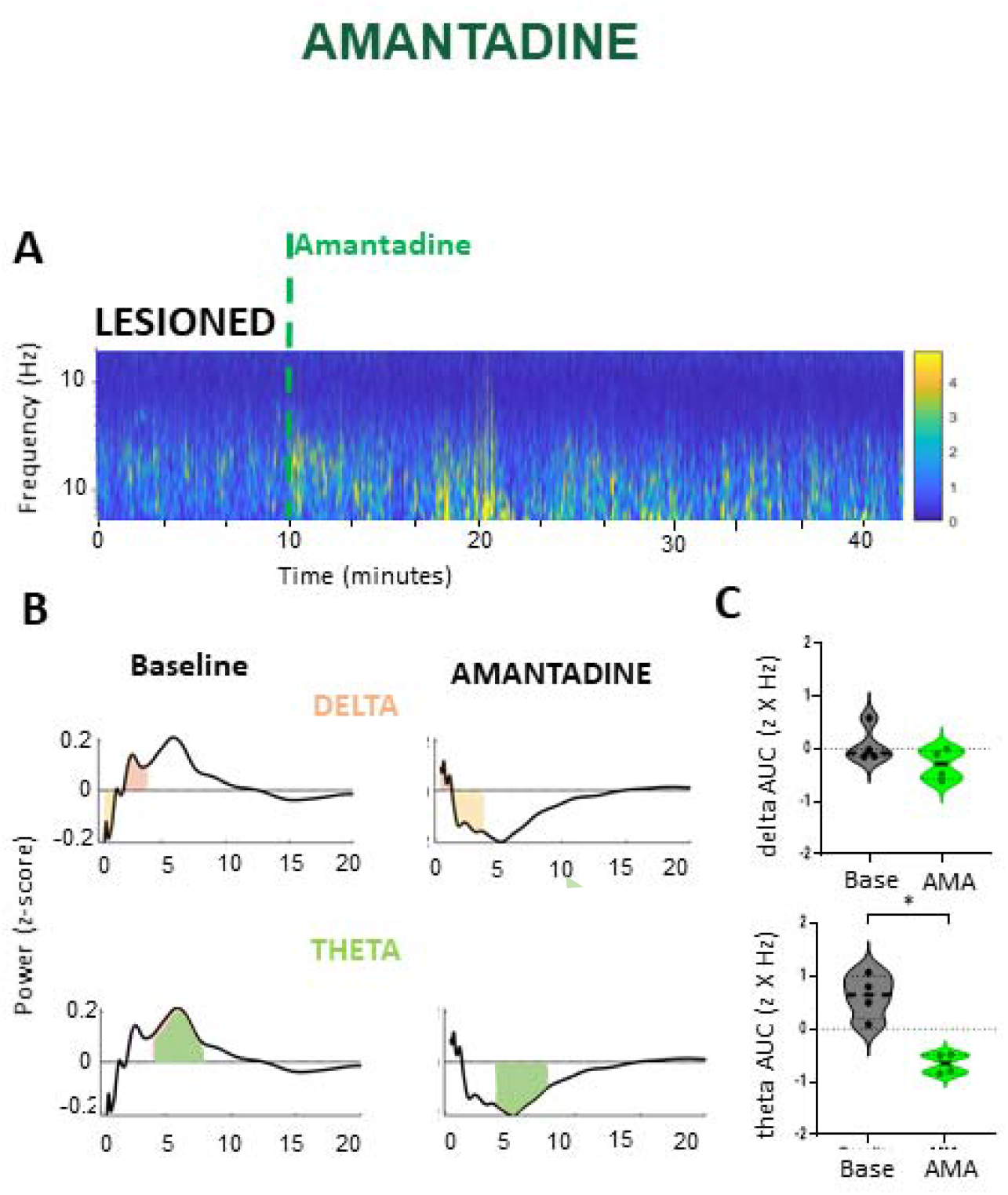
Amantadine reduces theta ACh oscillatory activity in DA-deprived state. **(A)** Average striatal ACh power spectral density is shown as colormap and **(B)** Z-scored power averaged over baseline time window (baseline: first 10 minutes) and in the presence of amantadine 60mg/kg (30-50 min from recording onset) is shown in frequency domain with AUC highlighted in the delta and theta bands. **(C)** Band-specific AUC analyses reveal a selective reduction of theta power in the presence of amantadine relative to baseline in DA-lesioned hemispheres (P=0.02), with no significant effect on delta oscillations (P=0.24, n=4, paired t-test)

## Notes

### Competing Interest Statement

The authors have declared no competing interest.

### Summary of Updates

This version of the manuscript has been revised to update the following: title, author's list, declaration of interest and acknowledgments.

